# Effects of Ambient Temperature During Pregnancy on Newborn Birthweight

**DOI:** 10.1101/2025.06.10.658978

**Authors:** Reshma Nargund, Joddy Marchesoni, Akshay Bareja, David W. Sosnowski, Gang Peng, Cathrine Hoyo, William Pan, Susan K. Murphy

**Affiliations:** Nicholas School of the Environment, Duke University, Durham, NC, 27708, United States; North Carolina State University, Raleigh, NC, 27695, United States; Department of Medicine, Duke University School of Medicine, Durham, NC 27701, United States; Duke Molecular Physiology Institute, Duke University, Durham, NC 27701, United States; Johns Hopkins Bloomberg School of Public Health, Baltimore, MD, 21205, United States; Department of Medical and Molecular Genetics, Indiana University School of Medicine, Indianapolis, IN, 46202, United States; Department of Obstetrics and Gynecology, Duke University, Durham, NC 27701, United States

## Abstract

This study evaluates the association between ambient temperature exposure during pregnancy and newborn birthweight, using a penalized generalized additive model (GAM) framework with distributed lag non-linear models (DLNM) to identify sensitive windows of exposure. The analysis includes 238 participants from the SHIP study with complete temperature exposure and birthweight data. Weekly maximum temperatures during pregnancy were estimated using Daymet data, and the impact of temperature on birthweight was assessed, adjusting for maternal age, pre-pregnancy BMI, gestational age, race, smoking, diabetes status, and infant biological sex. The model incorporated a crossbasis function for temperature exposure across 42 gestational weeks and allowed penalization for smoother, data-driven lag estimation. Results from the combined-sex model indicated that higher ambient temperatures during the third trimester, particularly in the final weeks of pregnancy, were associated with increased birthweight. Stratified analyses suggested that this association was more pronounced in male infants. These findings highlight the importance of considering prenatal temperature exposures and timing when evaluating determinants of newborn health.

## Introduction

### Background

Pregnant women and unborn children are increasingly being recognized as a population vulnerable to extreme temperature events (Spencer et al. 2022). Core body temperature (CBT) regulation in human adults is a delicate balance between environmental heat load, internal heat generation, and capacity for heat loss (Kurz 2008). CBT regulation in pregnancy faces additional challenges due to changes in cardiovascular, respiratory, gastrointestinal, and neurological systems, among others (Soma-Pillay et al. 2016). In addition to adverse physical health outcomes, extreme heat is also implicated in poor mental health and altered behavior in pregnant women (Lin et al. 2017). With the increase in average global temperatures, we can expect more frequent heat waves in addition to higher seasonal temperatures (Hoegh-Guldberg et al. 2019). Previous studies have examined associations between temperature and birthweight. A 2020 study in China of 4951 pregnant women showed that maternal exposure to 29 (95th percentile) degrees Celsius as compared to the reference temperature of 20 degrees Celsius was associated with a decrease in placental volume and weight, and an increase in the placental weight to birth weight ratio (PFR) (Wang et al. 2020). The placenta is the core interface between the mother and the fetus and performs critical physiological functions such as transportation of oxygen, nutrients, and waste (Philippat et al. 2019). Additionally, it is involved in the production, transfer, and metabolism of hormones critical to fetal development such as progesterone, estrogen, and human chorionic gonadotropin (hCG) (Murphy et al. 2006). Given the placenta’s critical role in fetal development, extremely low and high placental weights have been associated with adverse outcomes like pre-eclampsia (DAHLSTRØM et al. 2008), and low APGAR scores at birth (Eskild, Haavaldsen, and Vatten 2014). Given the placenta’s critical role in fetal development, it stands to reason that temperature-related changes in placental weight and the placental-to-birthweight ratio (PFR) could influence birthweight outcomes. Supporting this, a 2012 prospective study in 1,031 healthy singleton pregnancies found that placental weight is a significant determinant of fetal growth and birthweight, independent of maternal factors such as BMI, fasting glucose, and weight gain (Roland et al. 2012).

This study evaluated the association between temperature exposure during pregnancy and newborn birthweight using a penalized GAM framework with DLNM to identify critical windows of vulnerability. With this study, we asked whether ambient temperature during pregnancy influences newborn birthweight, and if so, which periods of gestation are most sensitive.

### Study Participants

Mother-infant dyads were enrolled from university-affiliated obstetric clinics between April 2018 and March 2020 as part of the Stress and Health in Pregnancy (SHIP) cohort based at North Carolina State University in Raleigh, North Carolina. Women were eligible if they were 18 years or older, spoke English or Spanish and planned to deliver at the study-affiliated hospital. Women were excluded from the study if the fetus had any known congenital anomalies or chromosomal disorders, or if the mother was diagnosed with HIV, Hepatitis C, or Hepatitis B. After enrollment, women completed questionnaires covering demographics, health behaviors, and provided obstetric, medical, and social histories. Maternal obstetric and infant delivery records were abstracted post-delivery. This study was approved by the institutional review board of North Carolina State University, and mothers provided written informed consent for both themselves and their children (Voegtline et al. 2025).

A total of 266 mother-infant dyads were included in the analysis. Of these, nine were excluded due to twin births, seven due to missing geocoded addresses, 12 due to missing delivery dates, four due to missing gestational age, one due to intersex birth, and one due to missing infant biological sex.

Twin births are typically excluded from birth weight outcome studies due to the association of multiple gestation pregnancies with intrauterine growth restriction and preterm birth (Garite et al. 2004). Participants with missing delivery dates (N=12), gestational age (N=4), or ungeocoded residential addresses (N=7) were excluded because accurate exposure assignment required complete temporal and spatial data. Specifically, we geocoded residential addresses and assigned temperature exposure values based on participants’ location at enrollment. For each participant, we used the date of birth and gestational age to calculate the pregnancy window—working backward from the date of delivery for the full gestational duration. Temperature values were then extracted for this pregnancy period. As such, participants missing delivery dates, gestational age, or a valid geocoded address could not be reliably assigned exposure data and were therefore excluded from the analysis.

Additionally, gestational age is a critical covariate in birth weight studies, as birth weight is closely linked to gestational duration. Participants with missing infant biological sex data (N=1) and intersex births (N=1) were excluded because infant biological sex is strongly associated with birth weight, and research suggests that male infants may be more susceptible to intrauterine conditions and environmental exposures (lampl et al. 2009). Mother’s age, BMI, smoking status, race and pre-pregnancy diabetes status were missing for 2, 2, 22, and 3 records, respectively. Race data were available for all participants. Race was categorized as Black, Hispanic, and White. Complete data were available for 238 participants.

### Ambient Temperature Exposure Assessment

Daily temperature exposure was estimated using the Daymet dataset, which provides high-resolution, gridded climate data across North America at a spatial resolution of 1 km x 1 km (Thornton et al. 2021). Maximum daily temperatures were assigned to participants based on geocoded residential addresses. The maximum daily temperature values were averaged to create weekly exposure estimates. Varying gestational lengths, which are a common challenge when dealing with time-series data and exposures during pregnancy were handled by implementing the Last Observation Carried Forward (LOCF) method. This approach ensures that participants are not excluded from the analysis due to missing exposure data for the weeks after birth (Steven J. Hadeed and Canales 2020). For participants with shorter gestational periods, the temperature value for the week of birth is carried forward to the remaining weeks. Specifically, for pregnancies shorter than 42 weeks, the temperature value for the week of birth was carried forward to assign a complete 42-week exposure profile based on residential location at enrollment. This ensures all participants have the same time series lenght.

### Statistical Analysis

We use a Distributed Lag Non-Linear Model (DLNM) within a penalized Generalized Additive Model (GAM) framework to evaluate the relationship between weekly temperature exposure during pregnancy and birthweight, adjusting for important covariates. The DLNM captures non-linear and delayed effects over time, which is particularly suitable for this type of study, as temperature exposure is expected to have non-linear effects that manifest over time. Specifically, lagged effects may result in a delayed impact of temperature on newborn birthweight, reflecting the complex, time-dependent nature of how environmental factors like temperature influence biological outcomes. Additionally, the DLNM allows for non-linear relationships between temperature exposure and outcome, accommodating the possibility that temperature effects may not be constant across all values. For instance, the impact of temperature could be more pronounced at extreme high or low temperatures. By modeling these effects, the DLNM provides flexibility in capturing both the delayed and level-dependent influence of temperature on newborn birthweight. On the other hand, the GAM framework allows for a flexible modeling approach for the relationship between predictors (such as temperature and maternal age) and the outcome, using smooth functions (e.g., splines). This flexibility is essential when the relationships between predictors and the outcome are not expected to be linear. The application of a penalty to the smooth functions helps prevent overfitting, ensuring that the model remains robust while capturing these complex relationships. Analysis was conducted in R using mgcv and dlnm (Gasparrini et al. 2017) that are designed for distributed lag non-linear penalized GAM.

Following the initial regression analysis conducted for the full sample, stratified analyses were performed for male and female infants, with separate models run for male and female infants. Additional stratified analyses were conducted based on maternal diabetes status (diabetic vs. non-diabetic) and infant growth classification (average for gestational age [AGA], small for gestational age [SGA], and large for gestational age [LGA]). Within the AGA subgroup, further stratification was performed by infant biological sex. Due to the limited sample size, stratified analyses by infant biological sex were not performed for the SGA and LGA groups.

## Results

### Descriptive Statistics

The mean age of the mothers was 29 years (SD = 6.1), with a median of 28.5 years, ranging from 19 to 43 years. The mean gestational age at birth was 38.5 weeks (SD = 1.8), with a median of 39 weeks, ranging from 30.3 to 41.6. Of the newborns, 132 were male (55.5%) and 106 were female (44.5%). The mean birth weight was 3208.9 grams (SD = 565.4 grams), with a median of 3227.5 grams (IQR = 2856.2–3558.8 grams), ranging from 1680 grams to 5000 grams. Among the newborns, 27 were classified as preterm (i.e., born before 37 weeks of gestation), representing 11.3% of the total births. Additionally, 36 newborns were classified as small for gestational age (SGA), 28 as large for gestational age (LGA), and 174 as average for gestational age (AGA), based on their birthweight percentile and gestational age at birth.

Distribution of key variables is shown in Table 1. Mean birth weight for males (3221.7 ± 615.5) was higher than females (3192.9 ± 498.4). There was no significant correlation between maternal age and birth weight (0.02, p = 0.82, 95% CI: -0.11 to 0.14) or pre-pregnancy BMI and birth weight (0.09, p = 0.15, 95% CI: -0.03 to 0.22). There was no significant correlation between pre-pregnancy diabetes status and birth weight (0.11, p= 0.09, 95% CI: -0.02 to 0.23), suggesting no strong association between these variables and birth weight. The distribution of birth weight was assessed for normality using skewness and kurtosis. Birth weight was approximately normally distributed with a skewness of -0.04 and kurtosis of 3.17.

**Table 1:**
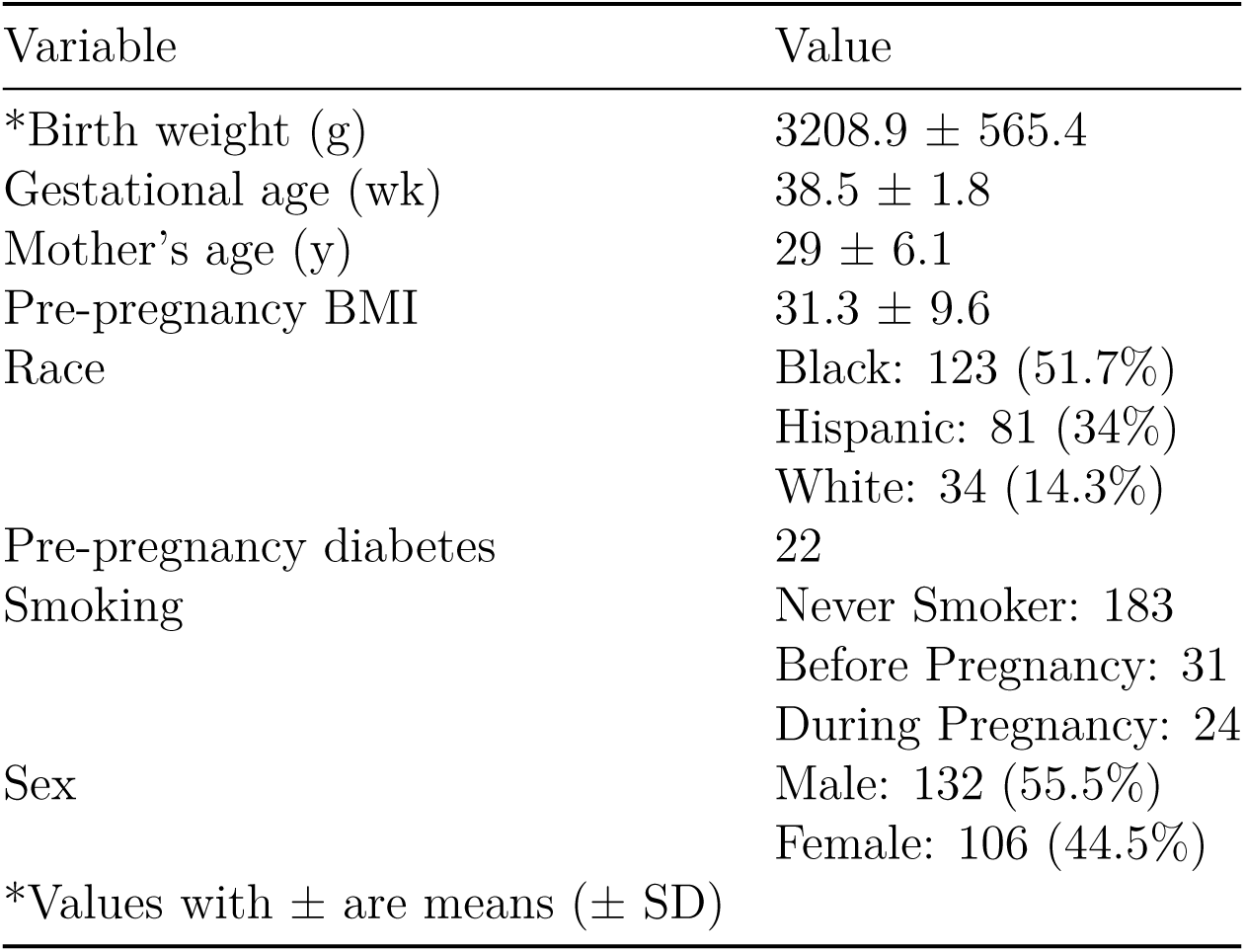
Key Variables Included in the Regression.

| Variable | Value |
| --- | --- |
| *Birth weight (g) | $3208.9 \pm 565.4$ |
| Gestational age (wk) | $38.5 \pm 1.8$ |
| Mother’s age (y) | $29 \pm 6.1$ |
| Pre-pregnancy BMI | $31.3 \pm 9.6$ |
| Race | Black: 123 (51.7%)<br>Hispanic: 81 (34%)<br>White: 34 (14.3%) |
| Pre-pregnancy diabetes | 22 |
| Smoking | Never Smoker: 183<br>Before Pregnancy: 31<br>During Pregnancy: 24 |
| Sex | Male: 132 (55.5%)<br>Female: 106 (44.5%) |
| *Values with ± are means (± SD) | |

Ambient temperature during pregnancy ranged from 0.8 °C to 37.2 °C, with an interquartile range of 16.8 °C to 32 °C. Temperature variation was observed across trimesters: in the first trimester (weeks 1–13), temperatures ranged from 0.8 °C to 37 °C; in the second trimester (weeks 14–26), from 0.8 °C to 37.2 °C; and in the third trimester (weeks 27–42), from 3.7 °C to 36.8 °C. A total of 188 participants experienced extreme weekly temperatures above 35 °C during their pregnancy, with these exposures most commonly occurring during the second trimester.

### Results of Penalized GAM DLNM (biological sexes combined)

A 1°C increase in ambient temperature during gestational weeks 36 to 40 was associated with a 19.1-gram increase in birth weight (95% CI: 4.65, 33.56; SE: 7.35) as shown in Figure 1. Full results of the regression analysis are presented in Table 2.

**Figure 1:**
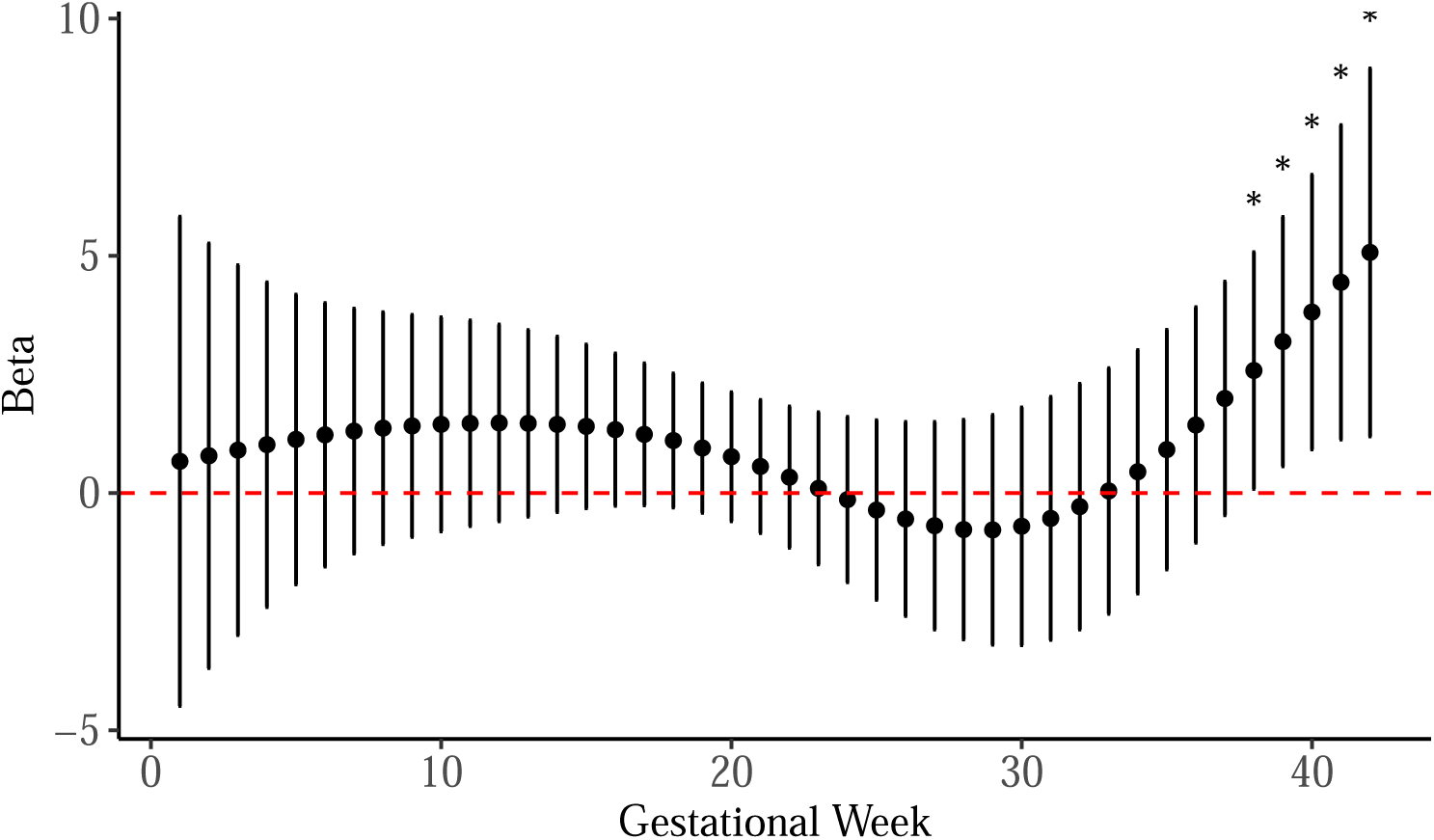
Weekly Gestational Temperature Effects on Birthweight - Biological Sexes Combined.

**Table 2:**
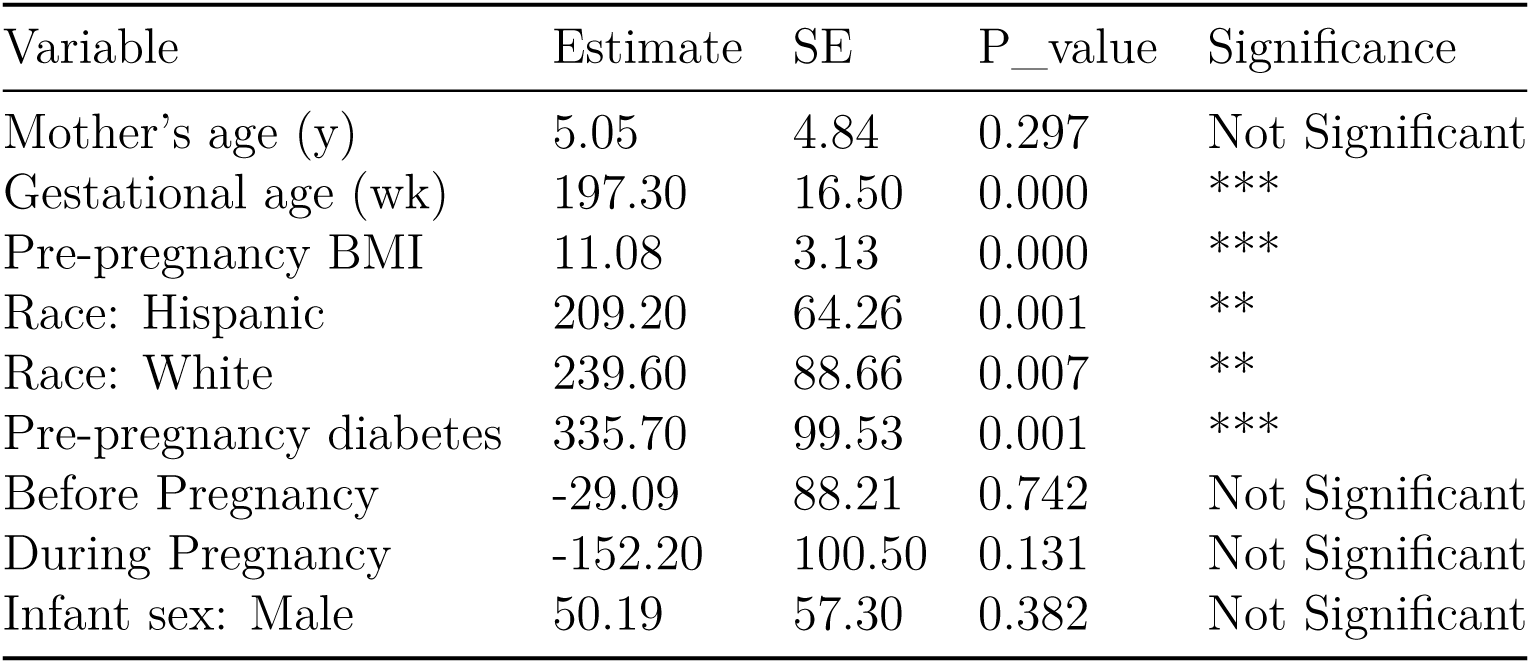
Regression Results - Biological Sexes Combined.

As expected, gestational age at birth was highly significant, with each additional week of gestation associated with a 197.3-gram increase in birth weight (95% CI: 165 - 229.68; p < 2 × 10 ¹). A one-unit increase in body mass index (BMI) at the last menstrual period corresponded to an 11.08-gram increase in birth weight (95% CI:4.95-17.2; p = 0.0004).

The overall contribution of race to the model was evaluated using an analysis of deviance comparing nested penalized GAMs. Inclusion of race significantly improved model fit (ΔDeviance = 3,144,339; df = 3.02; p = 0.0007), indicating that race was an important predictor of birthweight in the model.

Infants born to Hispanic mothers had, on average, a 209.2-gram higher birth weight compared to those born to Black mothers (95% CI: 83.25, 335.15; p = 0.001). Similarly, infants of White mothers had a 239.6-gram higher birth weight than those of Black mothers (95% CI: 65.83, 413.37; p = 0.007). Pre-pregnancy diabetes was also a significant factor, with affected mothers giving birth to infants who were, on average, 335.7 grams heavier than those born to mothers without diabetes (95% CI: 140.62, 530.78; p = 0.000877).

Male infants were, on average, 50.19 grams heavier than female infants; however, this difference was not statistically significant (95% CI: -62.12, 162.50; p = 0.382). The adjusted R² value for the model was 0.42, indicating that 42% of the variability in birth weight was explained by the included predictors. Maternal age and smoking status were not significant predictors of birth weight.

### Biological Sex-Specific Results of Penalized GAM DLNM

In males, a 1°C increase in ambient temperature during gestational weeks 35 to 40 was associated with a 39.5-gram increase in birth weight (95% CI: 14.52, 64.48; SE: 12.47). Similarly in males, a 1°C increase in ambient temperature during gestational weeks 8 to 17 was linked to a 27.8-gram increase in birth weight (95% CI: 5.75, 49.84; SE: 11.24) as shown in Figure 2.

**Figure 2:**
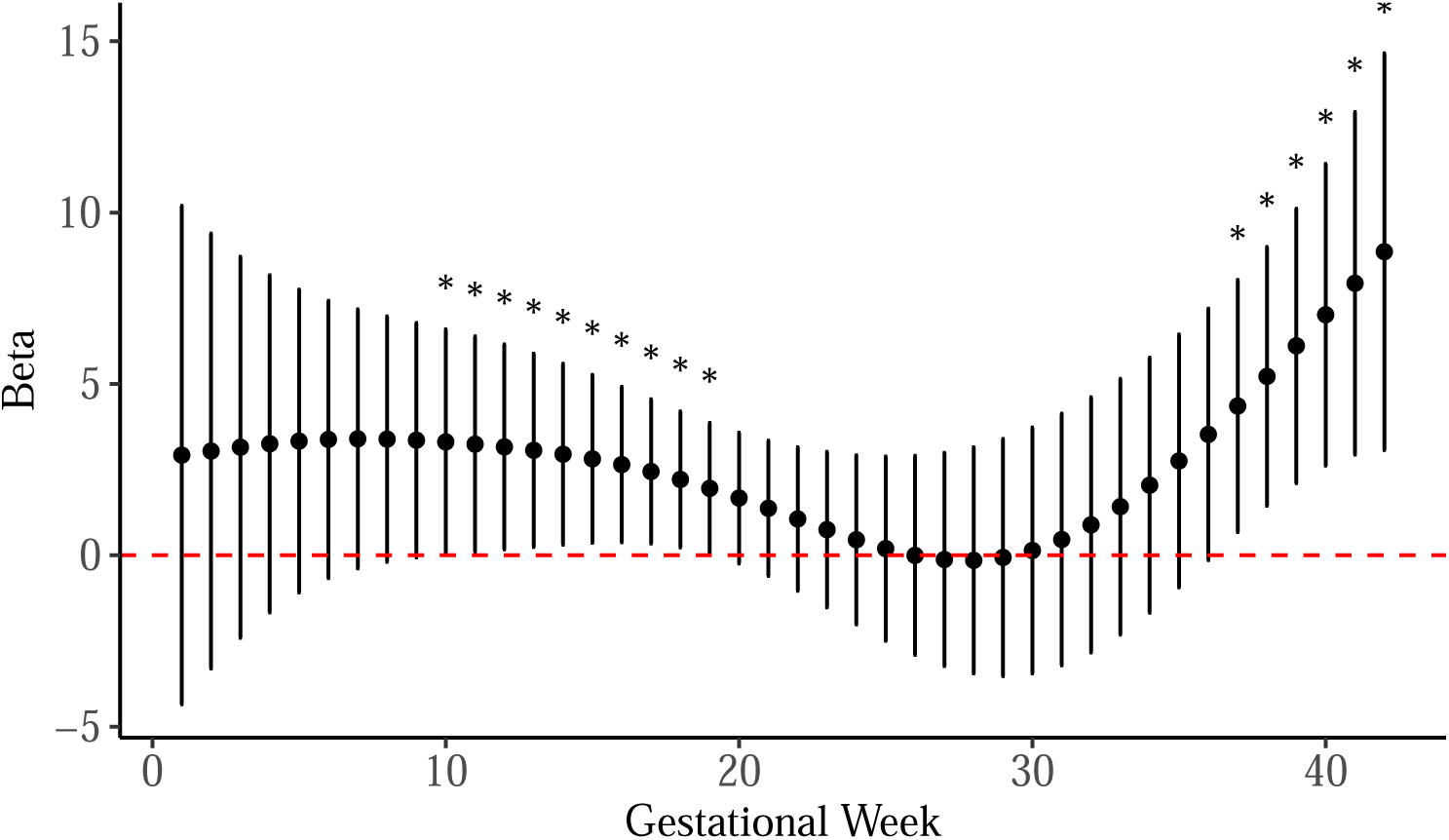
Weekly Gestational Temperature Effects on Birthweight - Male.

No effect of ambient prenatal temperature was seen in female infants (Figure 3). Key regression results for male and female infants are presented in Table 3 and Table 4.

**Figure 3:**
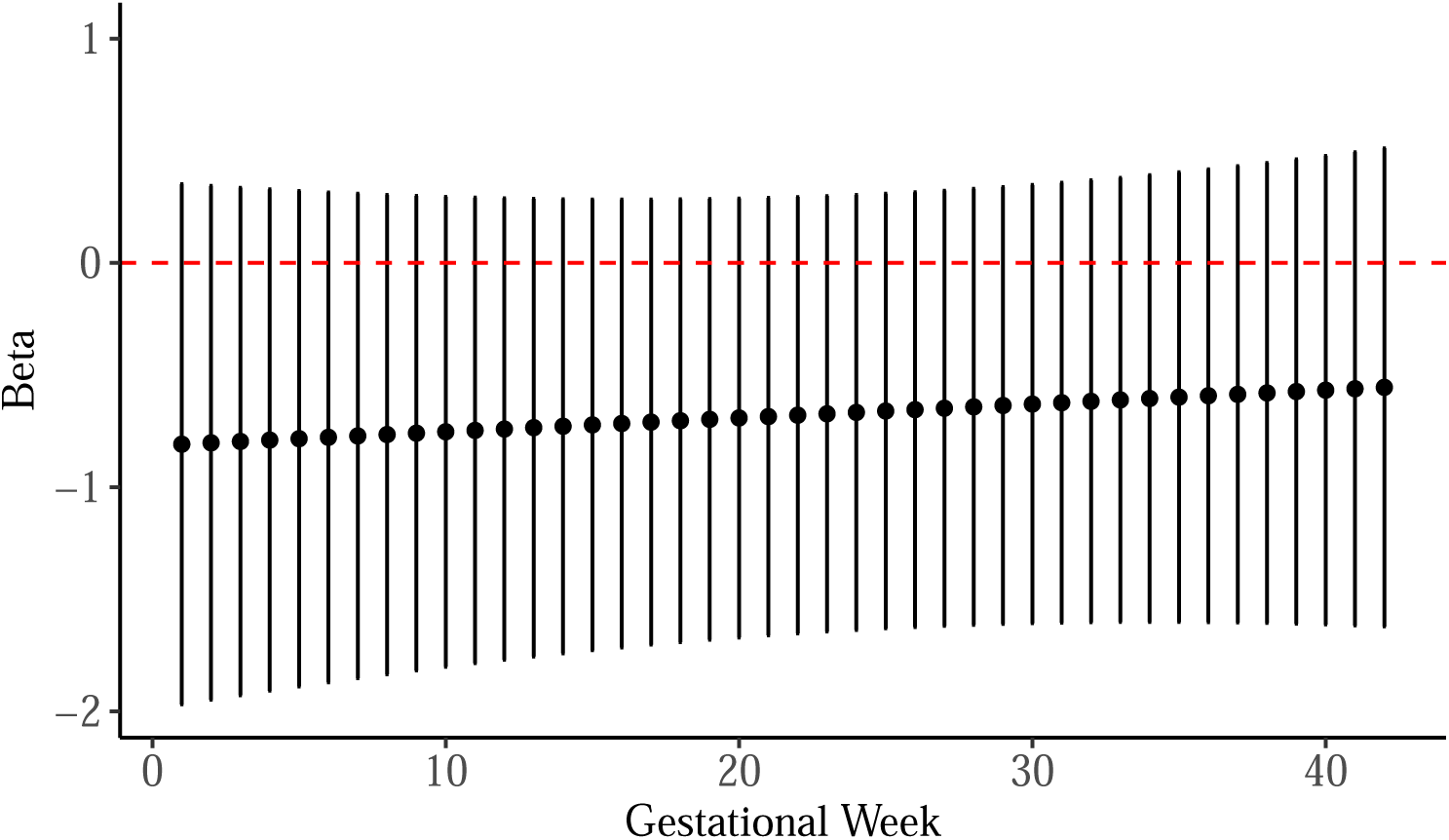
Weekly Gestational Temperature Effects on Birthweight - Female.

**Table 3:** Regression Results Males.

| Variable | Estimate | SE | P_value | Significance |
| --- | --- | --- | --- | --- |
| Mother's age (y) | 8.00 | 7.08 | 0.261 | Not Significant |
| Gestational age (wk) | 213.60 | 23.47 | 0.000 | *** |
| Pre-pregnancy BMI | 13.26 | 4.64 | 0.005 | ** |
| Race: Hispanic | 254.00 | 92.85 | 0.007 | ** |
| Race: White | 215.10 | 134.80 | 0.113 | Not Significant |
| Pre-pregnancy diabetes | 247.80 | 136.50 | 0.072 | . |
| Before Pregnancy | 8.17 | 126.90 | 0.949 | Not Significant |
| During Pregnancy | -241.10 | 164.90 | 0.146 | Not Significant |

**Table 4:** Regression Results Females.

| Variable | Estimate | SE | P_value | Significance |
| --- | --- | --- | --- | --- |
| Mother's age (y) | 5.05 | 6.69 | 0.452 | Not Significant |
| Gestational age (wk) | 187.98 | 23.82 | 0.000 | *** |
| Pre-pregnancy BMI | 6.68 | 4.32 | 0.125 | Not Significant |
| Race: Hispanic | 173.17 | 87.64 | 0.051 | . |
| Race: White | 231.47 | 122.22 | 0.061 | . |
| Pre-pregnancy diabetes | 475.55 | 155.05 | 0.003 | ** |
| Before Pregnancy | -147.49 | 121.11 | 0.226 | Not Significant |
| During Pregnancy | -77.50 | 121.33 | 0.525 | Not Significant |

### Stratified Analysis by Pre-pregnancy Diabetes Results of Penalized GAM DLNM

Mothers without pre-pregnancy diabetes exhibited no significant association between ambient temperature and infant birth weight (Figure 4). However, among infants born to mothers with pre-pregnancy diabetes, a 1°C increase in ambient temperature during gestational weeks 33 to 40 was associated with a 155.35-gram increase in birth weight (95% CI: 28.9, 281.8; SE: 64.5), as shown in Figure 5. Regression results are presented in Table 5 and Table 6.

**Figure 4:**
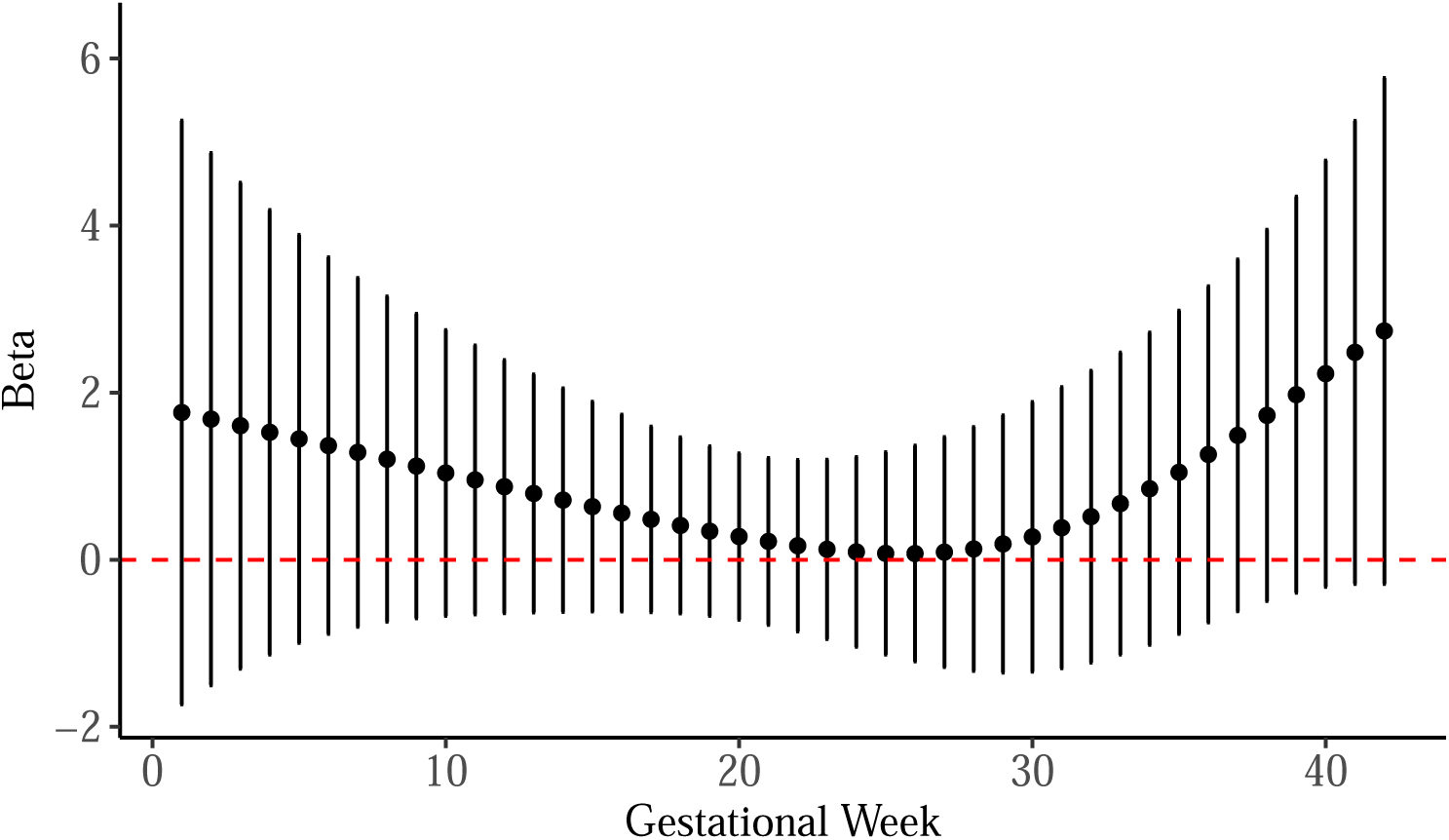
Weekly Gestational Temperature Effects on Birthweight - Non-Diabetic Mothers.

**Figure 5:**
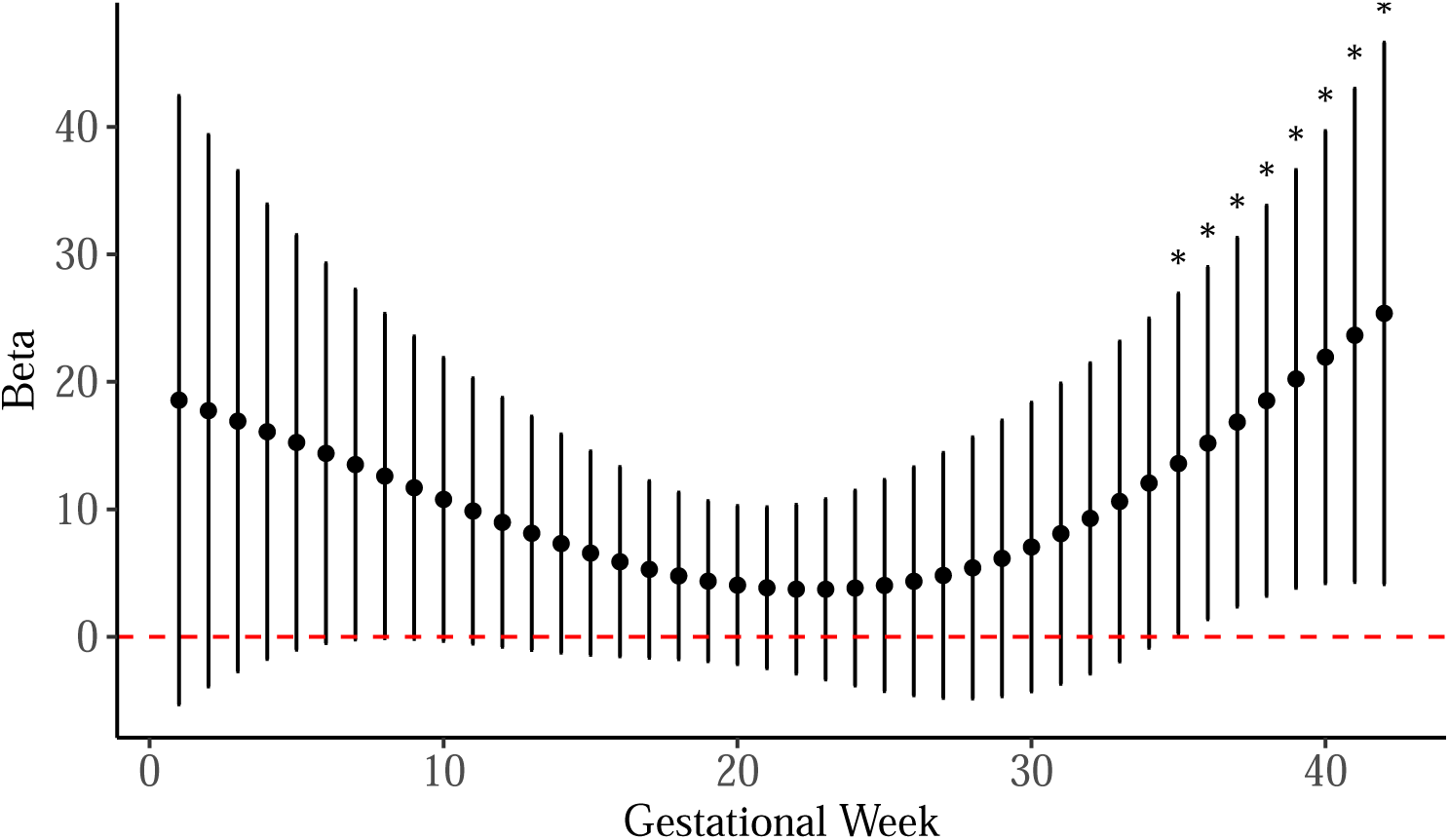
Weekly Gestational Temperature Effects on Birthweight - Diabetic Mothers.

**Table 5:** Regression Results - Non-Diabetic Mothers.

| Variable | Estimate | SE | P_value | Significance |
| --- | --- | --- | --- | --- |
| Mother's age (y) | 9.58 | 4.93 | 0.053 | . |
| Gestational age (wk) | 207.00 | 16.34 | 0.000 | *** |
| Pre-pregnancy BMI | 9.80 | 3.10 | 0.002 | ** |
| Race: Hispanic | 207.30 | 63.83 | 0.001 | ** |
| Race: White | 237.40 | 87.15 | 0.007 | ** |
| Before Pregnancy | -42.44 | 85.91 | 0.622 | Not Significant |
| During Pregnancy | -118.70 | 99.85 | 0.236 | Not Significant |
| Baby sex: Male | 87.95 | 57.66 | 0.129 | Not Significant |

**Table 6:** Weekly Gestational Temperature Effects on Birthweight - Diabetic Mothers.

| Variable | Estimate | SE | P_value | Significance |
| --- | --- | --- | --- | --- |
| Mother's age (y) | -25.02 | 27.65 | 0.387 | Not Significant |
| Gestational age (wk) | 84.78 | 121.41 | 0.501 | Not Significant |
| Pre-pregnancy BMI | 26.54 | 19.59 | 0.205 | Not Significant |
| Race: Hispanic | 52.35 | 414.61 | 0.902 | Not Significant |
| Race: White | 90.23 | 563.17 | 0.876 | Not Significant |
| Before Pregnancy | 588.44 | 992.43 | 0.566 | Not Significant |
| During Pregnancy | -367.76 | 621.97 | 0.568 | Not Significant |
| Baby sex: Male | -100.96 | 337.36 | 0.771 | Not Significant |

### Stratified Analysis by AGA, SGA, LGA Results of Penalized GAM DLNM

When participants were stratified by AGA, SGA, and LGA to evaluate potential differences in susceptibility to ambient temperature exposure during pregnancy, no significant association between temperature and birth weight was observed across gestational age-normalized weight groups (Figure 6, Figure 9, Figure 10). Regression results are shown in Table 7, Table 10, Table 11. Further stratification of AGA infants by biological infant sex revealed no significant effects (Figure 7, Figure 8). Regression results are shown in Table 8 and Table 9.

**Figure 6:**
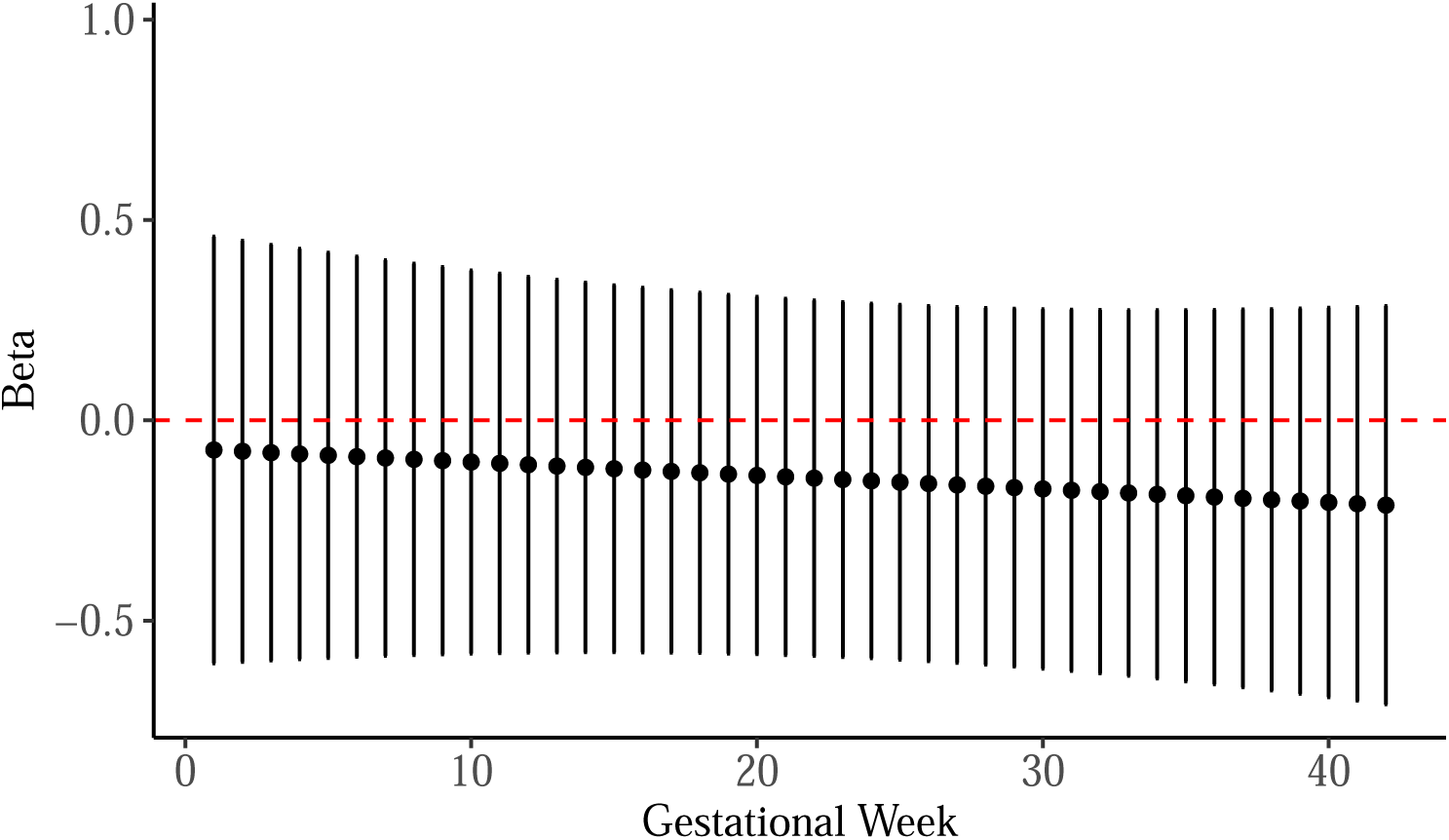
Weekly Gestational Temperature Effects on Birthweight - Average for Gestational Age.

**Figure 7:**
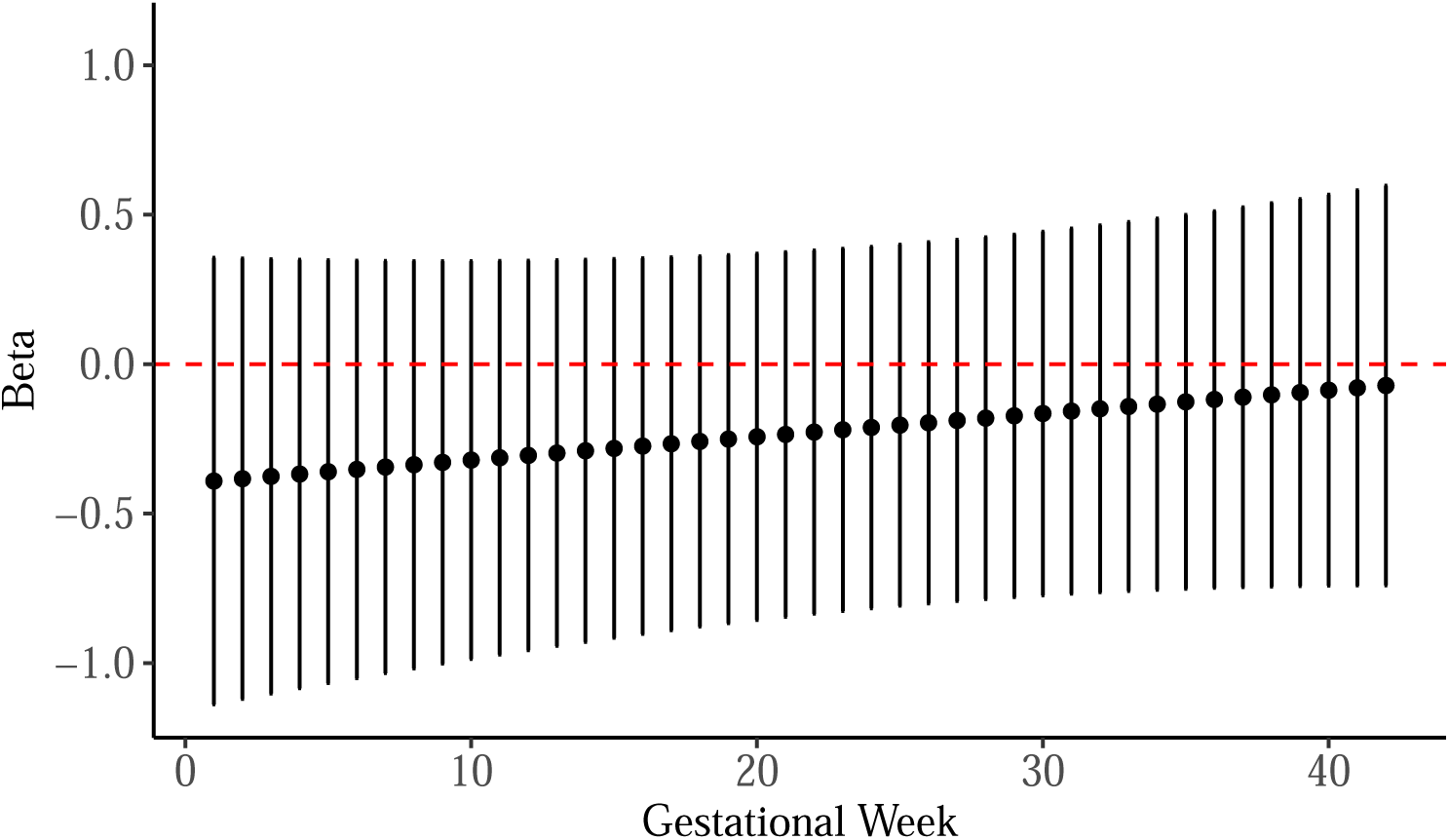
Weekly Gestational Temperature Effects on Birthweight - Average for Gestational Age - Male.

**Figure 8:**
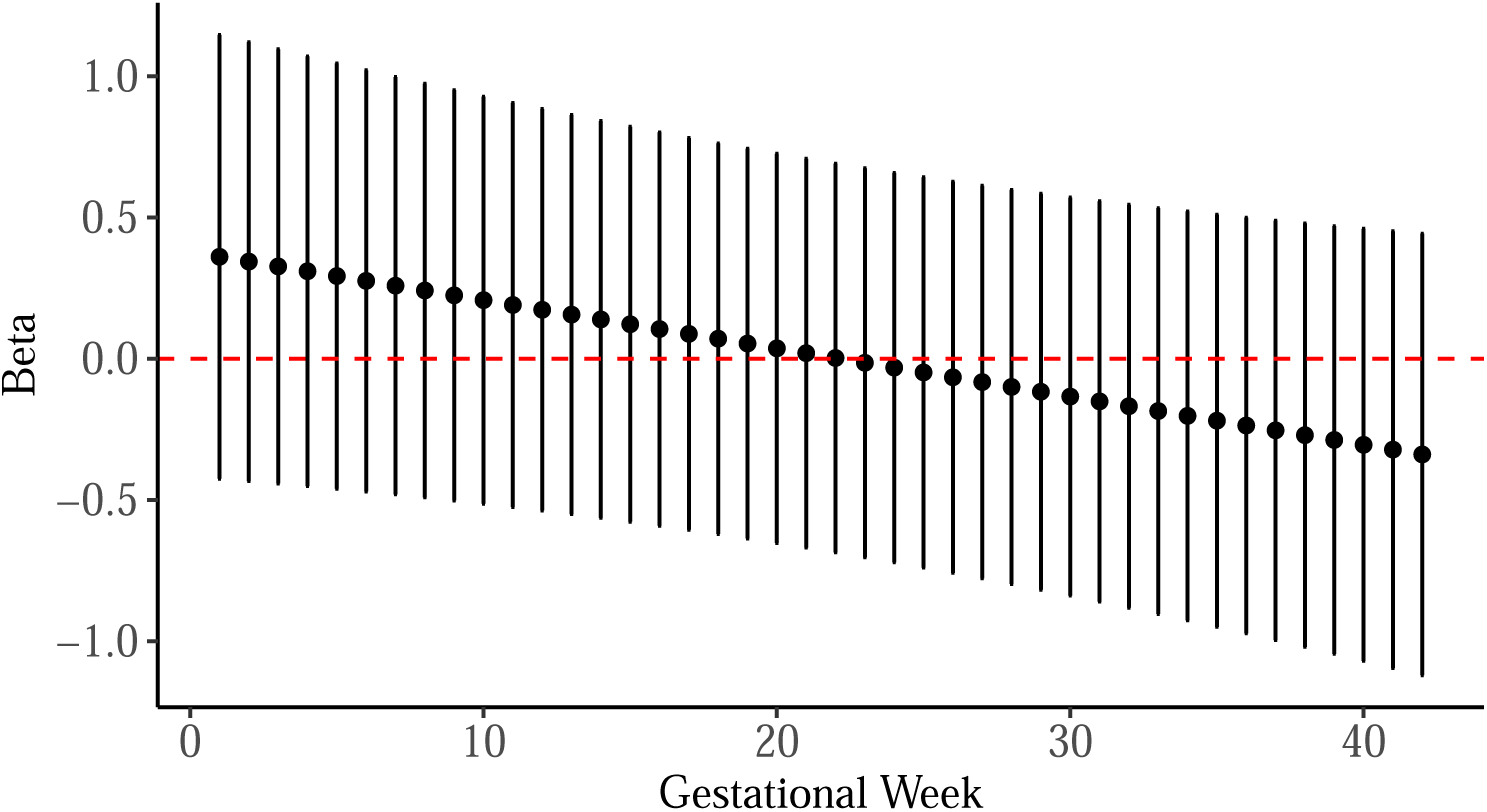
Weekly Gestational Temperature Effects on Birthweight - Average for Gestational Age - Female.

**Figure 9:**
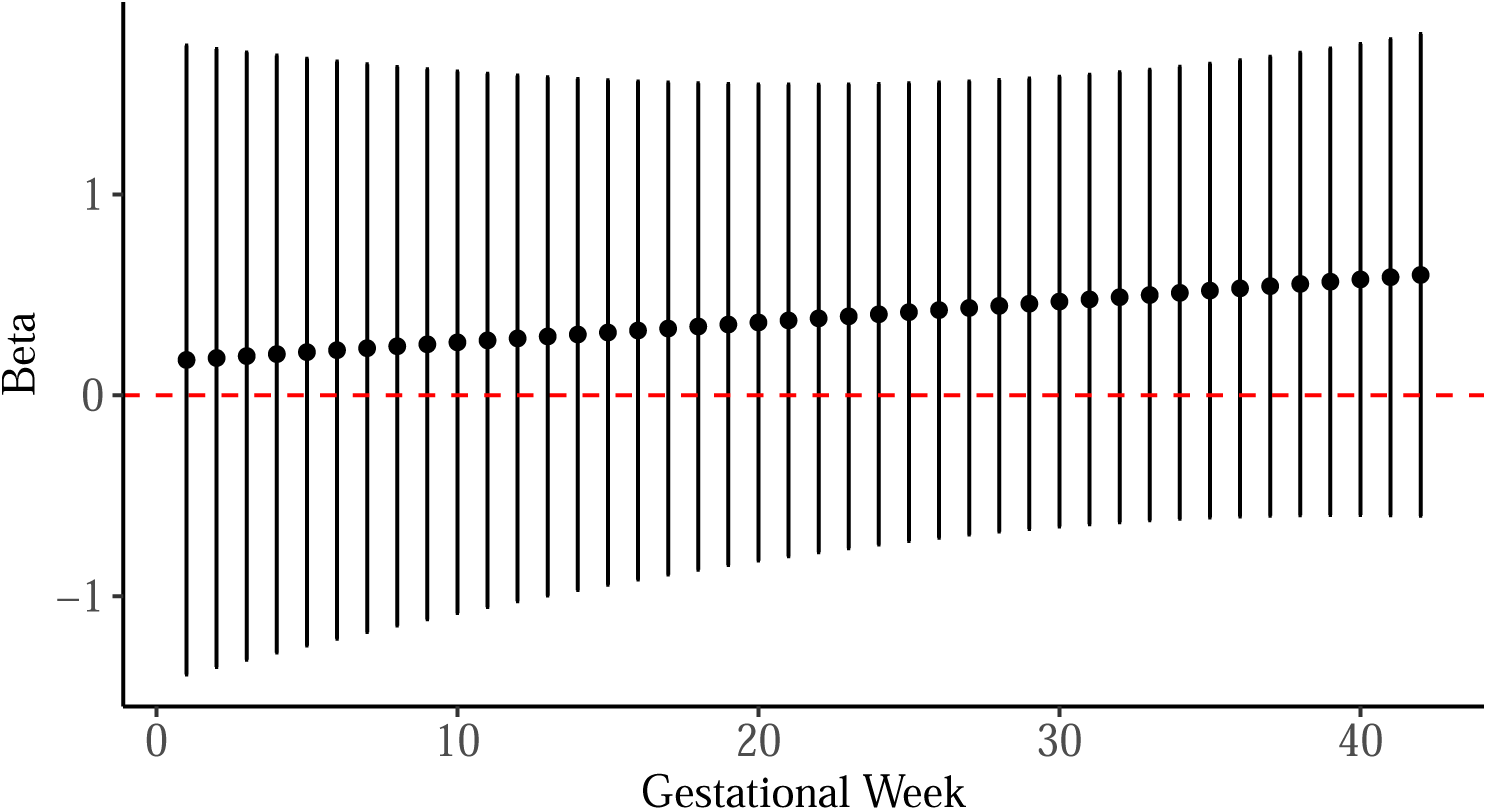
Weekly Gestational Temperature Effects on Birthweight - Small for Gestational Age.

**Figure 10:**
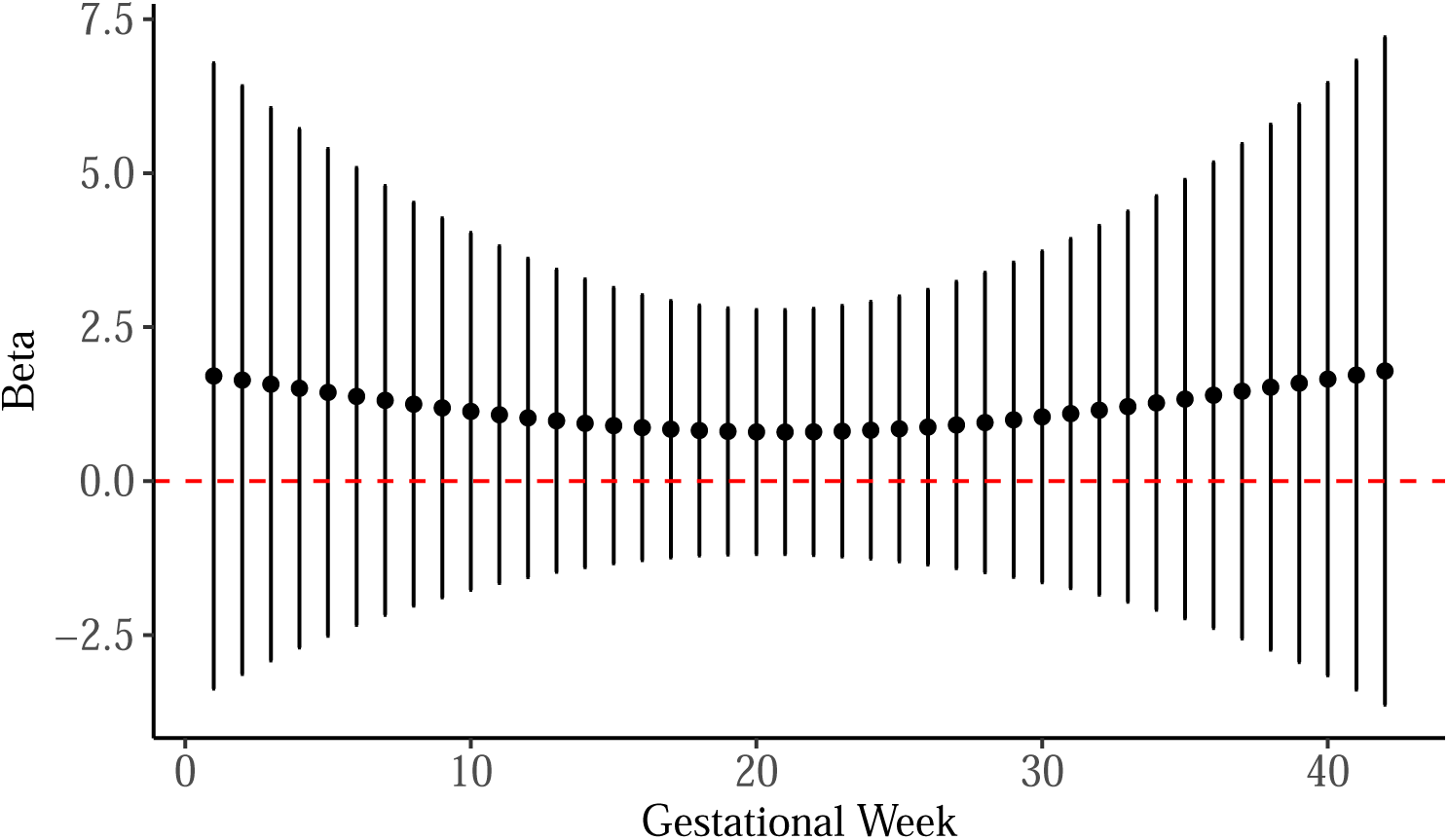
Weekly Gestational Temperature Effects on Birthweight - Large for Gestational Age.

**Table 7:**
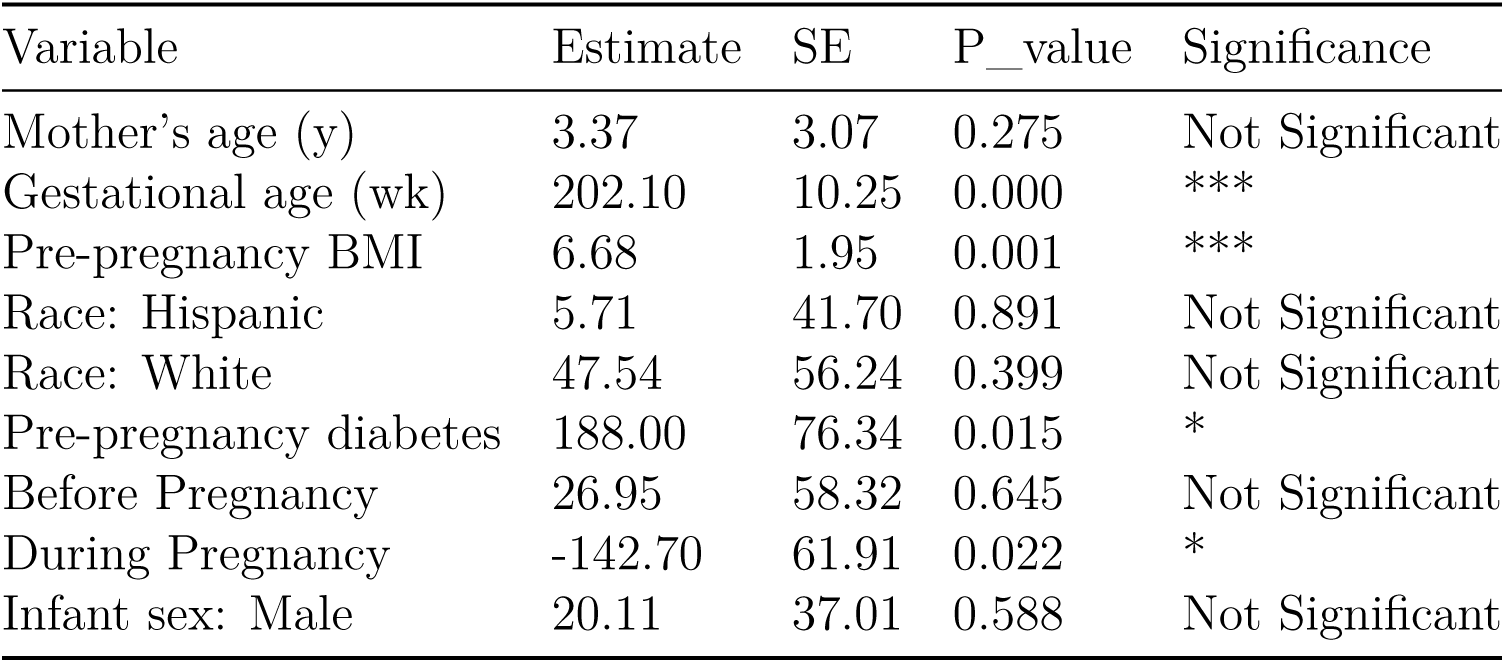
Regression Results - Average for Gestational Age.

**Table 8:** Regression Results - Average for Gestational Age – Male.

| Variable | Estimate | SE | P_value | Significance |
| --- | --- | --- | --- | --- |
| Mother's age (y) | 4.28 | 4.17 | 0.308 | Not Significant |
| Gestational age (wk) | 203.30 | 13.89 | 0.000 | *** |
| Pre-pregnancy BMI | 9.13 | 2.72 | 0.001 | ** |
| Race: Hispanic | 50.26 | 56.26 | 0.374 | Not Significant |
| Race: White | -15.60 | 80.31 | 0.847 | Not Significant |
| Pre-pregnancy diabetes | 249.40 | 90.18 | 0.007 | ** |
| Before Pregnancy | 130.80 | 81.98 | 0.115 | Not Significant |
| During Pregnancy | -164.80 | 91.83 | 0.077 | . |

**Table 9:** Regression Results - Average for Gestational Age – Female.

| Variable | Estimate | SE | P_value | Significance |
| --- | --- | --- | --- | --- |
| Mother's age (y) | 1.33 | 4.73 | 0.779 | Not Significant |
| Gestational age (wk) | 188.50 | 15.68 | 0.000 | *** |
| Pre-pregnancy BMI | 4.48 | 2.92 | 0.129 | Not Significant |
| Race: Hispanic | -33.28 | 62.45 | 0.596 | Not Significant |
| Race: White | 89.00 | 84.35 | 0.295 | Not Significant |
| Pre-pregnancy diabetes | 12.81 | 153.70 | 0.934 | Not Significant |
| Before Pregnancy | -43.18 | 83.53 | 0.607 | Not Significant |
| During Pregnancy | -148.40 | 84.91 | 0.085 | . |

**Table 10:** Regression Results - Small for Gestational Age.

| Variable | Estimate | SE | P_value | Significance |
| --- | --- | --- | --- | --- |
| Mother's age (y) | -20.21 | 8.58 | 0.027 | * |
| Gestational age (wk) | 132.68 | 34.60 | 0.001 | *** |
| Pre-pregnancy BMI | 1.79 | 4.95 | 0.720 | Not Significant |
| Race: Hispanic | 272.69 | 97.35 | 0.010 | ** |
| Race: White | -400.44 | 191.74 | 0.048 | * |
| Pre-pregnancy diabetes | -17.05 | 147.40 | 0.909 | Not Significant |
| Before Pregnancy | 114.06 | 107.11 | 0.298 | Not Significant |
| During Pregnancy | 411.39 | 184.94 | 0.036 | * |
| Infant sex: Male | -112.13 | 85.08 | 0.200 | Not Significant |

**Table 11:** Regression Results - Large for Gestational Age.

| Variable | Estimate | SE | P_value | Significance |
| --- | --- | --- | --- | --- |
| Mother's age (y) | -4.95 | 13.35 | 0.716 | Not Significant |
| Gestational age (wk) | 239.79 | 39.16 | 0.000 | *** |
| Pre-pregnancy BMI | 14.24 | 11.68 | 0.241 | Not Significant |
| Race: Hispanic | 289.93 | 189.35 | 0.145 | Not Significant |
| Race: White | 234.50 | 233.78 | 0.331 | Not Significant |
| Pre-pregnancy diabetes | 227.40 | 191.28 | 0.252 | Not Significant |
| Before Pregnancy | 416.28 | 258.03 | 0.126 | Not Significant |
| During Pregnancy | -269.60 | 377.11 | 0.485 | Not Significant |
| Infant sex: Male | 170.46 | 152.78 | 0.281 | Not Significant |

## Discussion

### Key Findings

Our results indicate that increased ambient temperature during late pregnancy is associated with higher birth weight. A 1°C rise in temperature during gestational weeks 36 to 40 corresponded to a modest increase in birth weight, with effect observed primarily in male infants during weeks 35 to 40. Earlier pregnancy exposure (weeks 8 to 17) was also linked to increased birth weight in males.

While the results of the stratified analyses summarized below are not statistically significant—because our approach involved running separate models rather than formally testing for interaction—they nevertheless point to potential effect modification by infant sex and maternal health status. Notably, among infants born to mothers with pre-pregnancy diabetes, the association was substantially greater, with late pregnancy temperature increases associated with a pronounced rise in birth weight. These findings suggest that fetal growth responses to ambient temperature may vary based on the biological sex of the infant and maternal health status, highlighting the need for further research into the underlying biological mechanisms and potential clinical implications. While we were not able to ascertain that males born to mothers with pre-pregnancy diabetes are most susceptible because of the small number of diabetic mothers, others have shown that males born to diabetic mothers are particularly susceptible to adverse birth outcomes (Maghalian, Alizadeh-Dibazari, and Mirghafourvand 2025).

### Biological Mechanisms

#### Third Trimester Sensitivity

A 2020 study demonstrated that fetal weight velocity, i.e., the rate of increase in weight over a specified time interval, peaked at week 35 of gestation, indicating that fetal weight gain accelerates in the third trimester (Grantz et al. 2018). This aligns with our finding that the effect of temperature on birth weight is most pronounced between weeks 35 and 40. Given that the rate of fetal weight gain peaks around week 35, it is possible that temperature exposure during this period accentuates the normal growth curve, leading to increased birth weight. This could be due to enhanced placental function or metabolic changes in response to temperature stress during this critical phase of rapid fetal development Wang et al. (2020). Additionally, pregnant women are often advised to increase their fluid intake during periods of high temperatures to mitigate the risk of dehydration. However, this increased need for hydration may inadvertently lead to higher consumption of sugared beverages, which could have implications for maternal and fetal health, particularly in terms of gestational diabetes and excessive weight gain (Chersich et al. 2020). Pregnant women may also do less physical activity during times of high temperature especially during the last weeks of pregnancy.

#### Infant Biological Sex Differences

Emerging evidence suggests that male fetuses exhibit greater vulnerability to prenatal stressors compared to females (Wu 2021), potentially due to sex-specific differences in placental function. Studies indicate that the male placenta shows distinct gene expression patterns, particularly in stress-responsive pathways, which may impair its ability to buffer maternal stress (Clifton 2010). For instance, research on asthma during pregnancy found that male fetuses responded by accelerating growth to enhance survival, whereas female fetuses reduced growth as a survival strategy (Clifton 2010). It is possible that this mechanism—accelerated fetal growth in response to stress in males—may also occur under other environmental stressors, such as exposure to elevated ambient temperatures. Hormonal differences, including testosterone surges in male pregnancies, may further exacerbate these vulnerabilities by altering placental signaling (Glezerman 2023). Collectively, these findings highlight how sex-biased biology contributes to differential outcomes in prenatal stress exposure.

#### Pre-pregnancy Diabetes and Temperature Sensitivity

Our findings suggest that infants of diabetic mothers exhibit a much stronger response to temperature, possibly due to pre-existing metabolic stress. Research has shown that maternal pre-pregnancy diabetes is linked to higher infant birth weight (Tyrrell et al. 2013). This suggests that one possible way temperature influences birth weight is by increasing the incidence and severity of maternal diabetes.

### Public Health and Climate Change Implications

Both being SGA and LGA can have perinatal, neonatal, and long-term health implications. Larger infants are susceptible to birth injuries such as shoulder dystocia, where the infant’s shoulder becomes stuck behind the mother’s pelvic bone (Øverland, Vatten, and Eskild 2013), clavicular fracture (Ahn et al. 2014), and Erb’s palsy, a type of brachial plexus injury (O’Berry et al. 2017). In addition to these birth injuries, high birth weight is linked to increased rates of cesarean deliveries (Poma 1999) and a higher risk of long-term health issues such as adolescent obesity (Zhao et al. 2012).

This study highlights the importance of tailoring maternal health advisories based on projected temperatures during pregnancy, particularly considering whether the third trimester will occur in the summer. Recommendations should account for the mother’s geographical location to ensure appropriate guidance. Beyond recommendations, it is essential to consider the cooling capacities of different communities and develop policies that address disparities in access to cooling resources. Mothers with pre-pregnancy or gestational diabetes, particularly those carrying boys, represent a highly vulnerable subgroup within an already at-risk population.

While pregnancy is a vulnerable period, the above findings—along with worsening temperature conditions due to climate change—highlight growing risks. Climate change is predicted to increase the frequency of extreme weather events, including cyclones, floods, wildfires, and more intense heat waves (Marx, Haunschild, and Bornmann 2021). These worsening temperature conditions further emphasize the need for tailored maternal health advisories and policies that address disparities in access to cooling resources, ensuring that vulnerable populations receive appropriate guidance and support.

### Strengths and Limitations

This study leverages advanced statistical methods, including distributed lag models, to capture the time-varying effects of environmental exposures during pregnancy. A key strength is the use of high-resolution temperature data, assigned based on geocoded residential addresses, allowing for detailed exposure assessment. Additionally, the incorporation of publicly available environmental datasets enhances transparency and reproducibility. However, some limitations must be acknowledged. Despite efforts to adjust for confounders, potential residual confounding remains a concern. Measurement error is also possible, particularly due to residential mobility, which may introduce exposure misclassification. Furthermore, while outdoor temperature exposure was assessed using residential locations, variations in indoor temperature, air conditioning use, and workplace exposures were not directly measured. Lastly, an imbalance in infant biological sex distribution could influence findings, necessitating careful interpretation of results.

### Future Directions

Future research should aim to further elucidate the biological mechanisms linking prenatal environmental exposures to infant health outcomes. If placenta samples were available, they could provide valuable insight into how these exposures influence the transfer of nutrients and fetal development at the molecular level. Additionally, integrating this work with epigenomic, transcriptomic and metabolomic data would allow for a more comprehensive understanding of exposure-related alterations in the placental epigenome, effects on placental gene expression and impacts on metabolic pathways. Such multi-omic approaches could help identify biomarkers of environmental exposure and potential mechanisms underlying observed associations, ultimately informing targeted interventions to improve maternal and child health outcomes.

## Conclusion

This study highlights the critical role of environmental exposures during pregnancy in shaping infant health outcomes. By leveraging high-resolution temperature data and advanced statistical methods, we observed significant associations between prenatal temperature exposure and birthweight. These findings underscore the importance of studying environmental influences during this sensitive developmental window, as exposure to extreme temperatures may have lasting effects on fetal growth trajectories and overall health. Given these implications, there is a pressing need for public health strategies aimed at mitigating the impact of extreme temperature events. Policies that promote access to cooling resources, improve housing conditions, and support for maternal adaptation to climate-related stressors could help protect vulnerable populations and promote healthier birth outcomes.

## Acknowledgements

This study was supported by NIH R01MD011746 (CH), Duke Global Health Doctoral Scholars Program, and von der Heyden Global Fellowship.

